# Cryo-EM structure of pro-aggregant P301L/S320F double-mutant tau filaments formed in mouse brains following peripheral AAV delivery

**DOI:** 10.64898/2026.01.19.700132

**Authors:** Maria Kano, Taeko Kimura, Haruaki Yanagisawa, Lisa Tatsumi, Komei Yamashita, Asahi Sakai, Xinyi Li, Takaomi C. Saido, Takashi Saito, Sho Takatori, Masahide Kikkawa, Taisuke Tomita

**Affiliations:** Laboratory of Neuropathology and Neuroscience, Graduate School of Pharmaceutical Sciences, The University of Tokyo, 7-3-1 Hongo, Bunkyo-ku, Tokyo 113-0033, Japan; Department of Cell Biology and Anatomy, Graduate School of Medicine, The University of Tokyo, 7-3-1 Hongo, Bunkyo-ku, Tokyo 113-0033, Japan; Laboratory for Proteolytic Neuroscience, RIKEN Center for Brain Science, 2-1 Hirosawa, Wako, Saitama 351-0198, Japan; Department of Neurocognitive Science, Institute of Brain Science, Nagoya City University Graduate School of Medical Science, 1 Kawasumi, Mizuho-cho, Mizuho-ku, Nagoya, Aichi 467-8601, Japan

**Keywords:** Alzheimer disease, cryo-EM, Pick disease tau, tauopathy

## Abstract

Tauopathies are characterized by the accumulation of abnormally phosphorylated tau filaments, with disease-specific folds revealed by cryo-electron microscopy (cryo-EM). Here, we delivered a recombinant human tau carrying the P301L/S320F double mutation into *App^NL-G-F^*/*MAPT* double knock-in mice using a blood-brain-barrier-permeable AAV (AAV-PHP.eB), enabling rapid induction of tau aggregation *in vivo*. The mutant tau was systemically administered via retro-orbital injection, providing a minimally invasive approach for widespread neuronal transduction. Phosphorylated tau aggregation, as well as seeding activity, were observed without the need for exogenous seeds. Our cryo-EM analysis resolved a novel filament fold incorporating residues 279-329, in which the P301L and S320F mutations introduced stabilizing interactions that reinforced filament assembly. This conformation was distinct from previously reported folds, including Alzheimer and Pick tau filaments. Moreover, the filaments adopted a more compact architecture than those observed in patient-derived samples or other model mice. This finding demonstrates that P301L/S320F double-mutant tau adopts a structurally unique, highly aggregation-prone fold, and that this system provides a rapid platform for modeling tau filament formation.

## Introduction

Tauopathies comprise a spectrum of neurodegenerative disorders defined by intracellular inclusions of abnormally phosphorylated and aggregated microtubule-associated protein tau, which is encoded by the Microtubule-Associated Protein Tau *(MAPT*) gene. These disorders include Alzheimer disease (AD), frontotemporal dementia (FTD), Pick’s disease (PiD), and corticobasal degeneration. In these conditions, the accumulation of tau filaments, known as neurofibrillary tangles (NFTs), is closely associated with progressive neurodegeneration and synaptic dysfunction [1,2].

Tau is an intrinsically disordered protein that stabilizes and dynamically regulates microtubules. The human *MAPT* gene generates six isoforms through alternative splicing, which differ in the number of microtubule-binding repeats (three-repeat (3R) or four-repeat (4R)) and the presence or absence of N-terminal inserts (Supplementary Fig. 1) [1]. In the adult human brain, 3R and 4R tau isoforms are expressed at approximately equal ratios, whereas murine brains predominantly express 4R tau [3]. The binding of tau to microtubules is tightly regulated by its phosphorylation state [4], and dysregulation of this balance is thought to underlie the onset of pathological aggregation. Notably, the discovery of pathogenic MAPT mutations in families with FTDP-17 provided strong genetic evidence that tau dysfunction is sufficient to cause neurodegeneration, and linked tau aggregation pathology to the pathogenesis of tauopathies [5,6].

In AD brains, aggregated tau forms paired helical filaments (PHFs) [7], which assemble into NFTs. This filament structure exhibits an AD-specific fold that reflects the disease’s molecular identity [8–10]. These filaments propagate in a prion-like manner between neurons, driving the stepwise regional spread of NFTs described by Braak staging [11]. The progression of NFT pathology correlates strongly with neuronal loss and cognitive decline [11,12], suggesting that tau filament accumulation directly drives neurodegeneration. Filamentous tau is not only hyperphosphorylated but also modified by ubiquitination [13–15], acetylation [16,17], and SUMOylation [18], among other post-translational modifications.

The advent of cryo-electron microscopy (cryo-EM) technology has enabled near-atomic structural determination of tau filaments isolated from tauopathy patient brains [8–10]. These structural studies have substantially advanced our understanding of the molecular mechanisms underlying tau aggregation and the pathological heterogeneity of tau-related disorders.

In parallel, *in vivo* modeling of tau pathology has advanced considerably. Transgenic mice expressing tauopathy-associated mutations [19] and models involving intracerebral inoculation of patient-derived [20] or recombinant tau filaments [21,22] have provided invaluable insights into the mechanisms of tau propagation and filament formation. More recently, expression of the P301L/S320F double-mutant tau— combining the frontotemporal dementia and Parkinsonism linked to chromosome 17 (FTDP-17)–associated P301L mutation [23,24] and the PiD-associated S320F mutation [25,26]—via adeno-associated virus (AAV) injection into neonatal mouse ventricles has been shown to induce robust tau aggregation within a short period [27]. This approach has attracted attention as a rapid and reproducible model of tau filament formation.

In this study, we sought to develop a less invasive and efficient *in vivo* model of AD-related tau pathology and to elucidate the structural determinants underlying the marked aggregation propensity of tau by analyzing the filamentous structures formed in this model. To this end, we utilized the AAV-PHP.eB system, a modified AAV9 variant that enables highly efficient central nervous system transduction following systemic administration, and expressed P301L/S320F mutant human tau *via* retro-orbital venous injection [28,29]. The AAV-PHP.eB serotype is reported to cross the blood–brain barrier following systemic administration efficiently and to robustly transduce cortical and hippocampal neurons, offering a less invasive alternative to direct intracranial injection. For the recipient mouse, we used humanized tau knock-in mice with an amyloid plaque– forming background (*App^NL-G-F^*/*MAPT* double knock-in mice), in which the murine *App* and *Mapt* genes are replaced by the human *APP* exons carrying familial AD-linked mutations as well as the *MAPT* locus, respectively [30]. This mouse model expresses both 3R and 4R human tau isoforms under endogenous control [31], with a comparable balance of tau isoforms to that observed in the human brain and also accumulates amyloid-β (Aβ) [30], thereby allowing investigation of tau aggregation within a genetic and pathological milieu that closely resembles the human AD brain.

Following systemic administration of AAV-PHP.eB encoding P301L/S320F mutant human tau, phosphorylated tau–positive aggregates were induced within approximately 1.5 months in a minimally invasive manner. Biochemical and cryo-EM analyses of sarkosyl-insoluble fractions revealed tau filaments with a near-atomic fold. These findings provide mechanistic insight into how the P301L/S320F mutation enhances tau aggregation propensity and support the emergence of a unique filament structure formed in a human tau background.

## Material and methods

### Mouse and cell culture

*App*^NL-G-F^/*hMAPT* double knock-in mice were maintained on a C57BL/6J background [32]. HEK293A cell was a kind gift from Dr. Hidenori Ichijo (The University of Tokyo)[33,34]. Biosensor (BS) cells, which provide a fluorescence resonance energy transfer (FRET) signal by seeding-competent tau filaments, were purchased from ATCC (CRL-3275). The cells were cultured as previously described [35,36].

### Extraction of the tau filaments from culture cells

Cells were resuspended in 1 mL of phosphate-buffered saline (PBS) and centrifuged at 4,500 × g for 5 min at 25 °C. The resulting pellets were resuspended in 300 µL of lysis buffer containing 1% sarkosyl in A68 buffer (10 mM Tris–HCl, pH 7.5, 10% sucrose, 0.8 M NaCl, and 1 mM EGTA) [37], followed by sonication using HG-30 homogenizer (Hitachi). An additional 300 µL of lysis buffer was then added, and the samples were centrifuged at 168,000 × g (S55A2-2698 roter, Himac) for 20 min at room temperature. The supernatant (soluble fraction) and pellet (insoluble fraction) were collected separately [38,39]. Protein concentrations in the supernatant were determined using a BCA assay, and 5× SDS sample buffer was added. The pellet fractions were resuspended in 50 µL of 2 × SDS sample buffer for western blotting, followed by brief sonication. All samples were heated at 100 °C for 5 min prior to electrophoresis. Prepared samples were separated by SDS–PAGE and transferred onto polyvinylidene difluoride (PVDF) membranes (Millipore) as described[35]. The list of primary antibodies and their dilution ratios is provided in Tables S1 and S2.

### Fluorescence resonance energy transfer (FRET)

BS cells were seeded into 24-well plates, and 20 µg of mouse brain homogenate was added to each well. The homogenate was mixed with Lipofectamine 2000 (Thermo Fisher Scientific) at 1.47 µL per well, incubated for 15 min at room temperature, and then applied to the cells. After treatment, the cells were cultured for 72 h. After incubation, cells were washed with PBS, detached by trypsinization, and resuspended in PBS containing 20% FBS. FRET signals were measured by flow cytometry using the MA900 cell sorter (SONY), with the increase in acceptor fluorescence upon donor excitation as the readout. The percentage of FRET-positive cells was calculated as described [40].

### Electron microscopy

For electron microscopy of cultured cell-derived tau filaments, pellet fractions were resuspended in 50 µL of Tris buffer (10 mM Tris-HCl, pH 7.4, 1 mM DTT, 1 mM EGTA). After sarkosyl fractionation of brain homogenates, the sarkosyl-insoluble pellet was resuspended in 10 µL of physiological saline, and 1.5 µL was applied onto collodion-coated 200 mesh copper grids (Nisshin EM Co). Negative staining was performed with 2% phosphotungstic acid. Samples were examined using a transmission electron microscope (JEM-1200EXII, JEOL) operated at an acceleration voltage of 80 kV. Images were acquired at magnifications ranging from 12,000× to 200,000× using a CCD camera (Veleta, JEOL) controlled with iTEM software (Olympus Soft Imaging Solutions). For immuno-electron microscopy, purified tau filaments were adsorbed onto collodion-coated 200 mesh copper grids (Nisshin EM Co.) for 30 s, followed by blocking with PBS containing 1% gelatin for 10 min and incubation with primary antibodies for 1 h. The grids were then washed with blocking buffer and incubated with 10 nm gold-conjugated anti-rabbit IgG or 5 nm gold-conjugated anti-mouse IgG (1:50 in blocking buffer) for 1 h (Table S1, 2). After washing with water, the samples were negatively stained with 2% PTA for 90 s[41].

### Recombinant AAV system

A recombinant AAV system of the PHP.eB [42] serotype was used to express human 1N4R MAPT carrying the P301L and S320F mutations. The expression cassette consists of the human MAPT cDNA with the P301L/S320F mutation under the control of the human synapsin I promoter, followed by a woodchuck hepatitis virus post-transcriptional regulatory element (WPRE) and a polyadenylation signal. The expression cassette was inserted into the pAAV-hSyn1-eYFP backbone (Addgene plasmid #117382) [29]. The eYFP coding sequence was removed by digestion with EcoRI and KpnI, and the MAPT insert was introduced using HiFi DNA Assembly (New England Biolabs). The recombinant AAV viruses used for mouse injections were produced and purified by the Department of Neurodegenerative Medicine, Gunma University Graduate School of Medicine. Recombinant virus production was carried out in HEK293T cells using the triple plasmids; pAAV-hSyn1-tau-WPRE, pHelper (Agilent Technologies), pAAV-PHP.eB [43,44]. AAV-PHP.eB were produced using the ultracentrifugation method described in a previous paper[43]. For *in vivo* experiments, recombinant viruses were delivered to mice via the retro-orbital sinus at a titer of 2–5 × 10¹¹ vg per mouse.

### Immunohistochemistry and image quantification

Immunohistochemical analyses were performed as previously described [45]. Stained sections were examined using fluorescence microscopy (BZ-X800, Keyence). High-resolution tiled images of AT8, AT180, PHF-1, and pS396 were reconstructed by stitching without compression, and exported using the Keyence analyzer package (BZ-X series). For the quantitative analysis of protein associations with phospho-tau lesions, we used a 20 × objective lens. Regions of interest were manually selected and cropped from the acquired images based on anatomical landmarks. The percentage of areas stained by AT8, AT180, PHF-1, and pS396 in selected brain regions was quantified using the BZ-X800 Analyzer Hybrid Cell Count Module (Keyence). Parameters were set to a threshold of control and applied automatically to all images within the proper comparisons. To minimize the influence of background staining and potential artifacts, AT8-or pS396-positive regions exceeding 10,000 µm² in area were excluded from the quantification. This threshold was set to filter out abnormally large signals likely due to non-specific staining. Statistical analyses were conducted using R software (version 4.5.1). Group means were compared using one-way analysis of variance (ANOVA), followed by Tukey’s Honest Significant Difference (HSD) test for post-hoc analysis (\**p* < 0.05).

### Sampling of mouse brains for western blotting

Fractionation of mouse brains for western blotting was performed as previously described with some modifications [46]. Half brains were homogenized in 10 volumes of homogenization buffer (10 mM Tris-HCl, pH 7.4, 1 mM EGTA, 1 mM EDTA, 0.8 M NaCl, 10% sucrose, supplemented with PhosSTOP (Sigma) and a protease inhibitor cocktail (cOmplete™ ULTRA, 1 tablet/50 mL; Roche) using a HG-30 homogenizer. The homogenates were ultracentrifuged at 168,000 × g for 20 min at 25 °C with an S55A2-2698 rotor. Sarkosyl was then added to the supernatant at a final concentration of 2%, followed by shaking at 37 °C for 30 min. After sonication, the samples were centrifuged at 20,000 × g for 20 min at 25 °C with an S55A2-2698 rotor, and the resulting supernatant was further ultracentrifuged at 168,000 × g for 20 min at 25 °C using an S55A2-2698 rotor. The pellet was resuspended in homogenization buffer containing 1% sarkosyl and subjected to the same centrifugation cycle. The final pellet was resuspended in Tris buffer, mixed with an equal volume of 2× SDS sample buffer, and boiled at 100 °C for 10 min to obtain the sarkosyl-insoluble fraction. Supernatants obtained at each centrifugation step were combined with sample buffer and used as sarkosyl-soluble fractions.

### Extraction of the tau filaments from mouse brains

Mouse brains were weighed and homogenized in 10 volumes (w/v) of homogenization buffer. Homogenization was performed using the HG-30 homogenizer (Hitachi). The homogenates were centrifuged at 11,400 × g (SW41Ti rotor, Beckman Coulter) for 10 min at room temperature, and the supernatants were collected. The supernatants were then centrifuged at 183,000 × g (SW41Ti rotor) for 4 h at 25 °C. The resulting pellets were resuspended in 1% sarkosyl solution prepared with half the volume of the original tissue buffer and incubated at 37 °C for 60 min with shaking. After incubation, the suspensions were centrifuged at 12,100 × g for 20 min at 25 °C (S55A2-2698 rotor), and the supernatants were collected. These supernatants were further centrifuged at 168,000 × g for 20 min at 25 °C (S55A2-2698 rotor), and the resulting pellets were resuspended in 10 mL of tissue buffer. To remove contaminating components, the suspension was treated with 0.2 mg/mL collagenase (Sigma-Aldrich), 2 mM MgCl₂, 0.5 mM CaCl₂, and 200 U/mL DNase I (Takara), followed by shaking at 200 rpm at 37 °C for 1 h. After enzymatic digestion, the final concentration of NaCl was adjusted to 1 M, and Triton X-100 was added to 0.5% (v/v). The mixture was further incubated at 37 °C for 30 min with shaking. To precipitate proteins, three volumes of pre-chilled methanol at –30 °C were added, and the samples were incubated for 30 min at –30 °C, followed by centrifugation at 20,000 × g for 30 min at 4 °C. The resulting pellets were resuspended in 10 mL of 5% sarkosyl solution. Subsequently, the following fractionation steps were repeated: centrifugation at 11,400 × g for 20 min at 25 °C to remove debris, followed by centrifugation of the supernatants at 168,000 × g for 30 min at 25 °C (S55A2-2698 rotor). The resulting pellets were resuspended in 2 mL of 5% sarkosyl solution, homogenized by passing through a 25-gauge needle (Terumo), and sequentially treated with 2% and 1% sarkosyl solutions in the same manner. At each step, the suspensions were mixed thoroughly by pipetting (100 times) before centrifugation. Finally, the pellets obtained were resuspended in 500 µL of 1% sarkosyl solution, homogenized again by pipetting, and centrifuged at 168,000 × g for 30 min (S55A2-2698 rotor). The final pellets, containing the sarkosyl-insoluble tau filaments, were collected and stored at –80 °C until use.

### Cryo-Electron Microscopy (cryo-EM)

Samples were resuspended in cryo-EM buffer (50 mM Tris-HCl (pH 7.4), 1 mM DTT, 0.04% sarkosyl [47]). A 2 µl aliquot of the sample was applied to each side of freshly glow-discharged UltrAuFoil R1.2/1.3 300 mesh grids (Quantifoil Micro Tools GmbH). The grids were blotted from both sides for 4 seconds at 12°C under 100% humidity and subsequently plunge-frozen in liquid ethane using a Vitrobot Mark IV (Thermo Fisher Scientific).

Images were recorded on a CRYO ARM 300 II (JEOL, Tokyo, Japan), operated at 300 kV, equipped with an Omega filter whose slit width was set to 20 eV, and a Gatan K3 direct electron detector in correlated-double sampling (CDS) mode. The nominal magnification was set to 60,000×, yielding a physical pixel size of 0.878 Å/pixel. Movies were acquired using the SerialEM software [48], with a target defocus range of −0.8∼-1.8 µm. Each movie was recorded with a total electron dose of 50 e⁻/ Å^2^, divided into 50 frames.

### Data processing of Cyro-EM data

Raw movie frames were motion-corrected, and the contrast transfer function (CTF) was estimated using the Patch CTF function on cryoSPARC v4.7.1[49]. Fibrils were automatically picked using the filament tracer tool, and segments were extracted with a box size of 420 pixels scaled to 300 pixels and an inter-box distance set to 10% of the box size. Several rounds of 2D classification were performed to exclude bad classes that lacked discernible helical twists and non-amyloid contaminants. Initial helical parameters were estimated from 2D class averages with a box size of 1000 pixels scaled to 750 pixels, and an initial reference was generated using the relion_helix_inimodel2d program [50]. The helical refinement was performed using local helical parameter searches in cryoSPARC. After further local and global CTF refinement, reference-based motion correction, and 3D classification, the particle coordinates were converted into Relion STAR format using UCSF pyem [51] with in-house modifications to convert metadata generated by the filament tracer tool to Relion’s counterparts. Relion v5.0 [52,53] was used for further CTF refinement and the 3D auto-refine with local helical parameter searches.

### Model building and refinement

An initial model was generated using ModelAngelo[53], and the model was refined using Coot v0.9.8.1 [54], Phenix v1.21.2-5419 [55], and Servalcat v0.4.105 [56]. The refined models were validated using Phenix. Image rendering was performed using UCSF ChimeraX [57] and MaskChains tool [58]. The schematic figures were produced using atom2svg.py [59].

## Result

### P301L/S320F double-mutant tau induced tau aggregation in cultured cells

In the human brain, six tau isoforms are generated by combinations of the number of microtubule-binding repeats,3R or 4R, and the presence of 0, 1, or 2 N-terminal insertions (Fig. S1).To partially recapitulate this isoform expression of the 3R and 4R pattern, we generated HEK293A-derived cell lines that stably express the representative 3R and 4R tau isoforms, 1N3R and 1N4R human tau. We then transfected these cells with a plasmid encoding the 1N4R P301L/S320F mutant tau (Fig. 1a). Three days after transfection, cells were collected and separated into sarkosyl-soluble and - insoluble fractions (Fig. 1b). Western blot analysis showed that P301L/S320F double-mutant tau was prominently detected in the sarkosyl-insoluble fraction, exhibiting a higher signal compared with wild-type (WT) tau. This finding indicates that the P301L/S320F mutation promoted the tau aggregation in cells, as previously described (Fig. 1c, d). Furthermore, electron microscopy of the sarkosyl-insoluble fraction revealed fibrillar structures with a twisted morphology and a diameter of approximately 20 nm (Fig. 1e). Taken together, these findings suggest that P301L/S320F tau can promote highly intracellular aggregation and filament formation.

**Figure 1.**
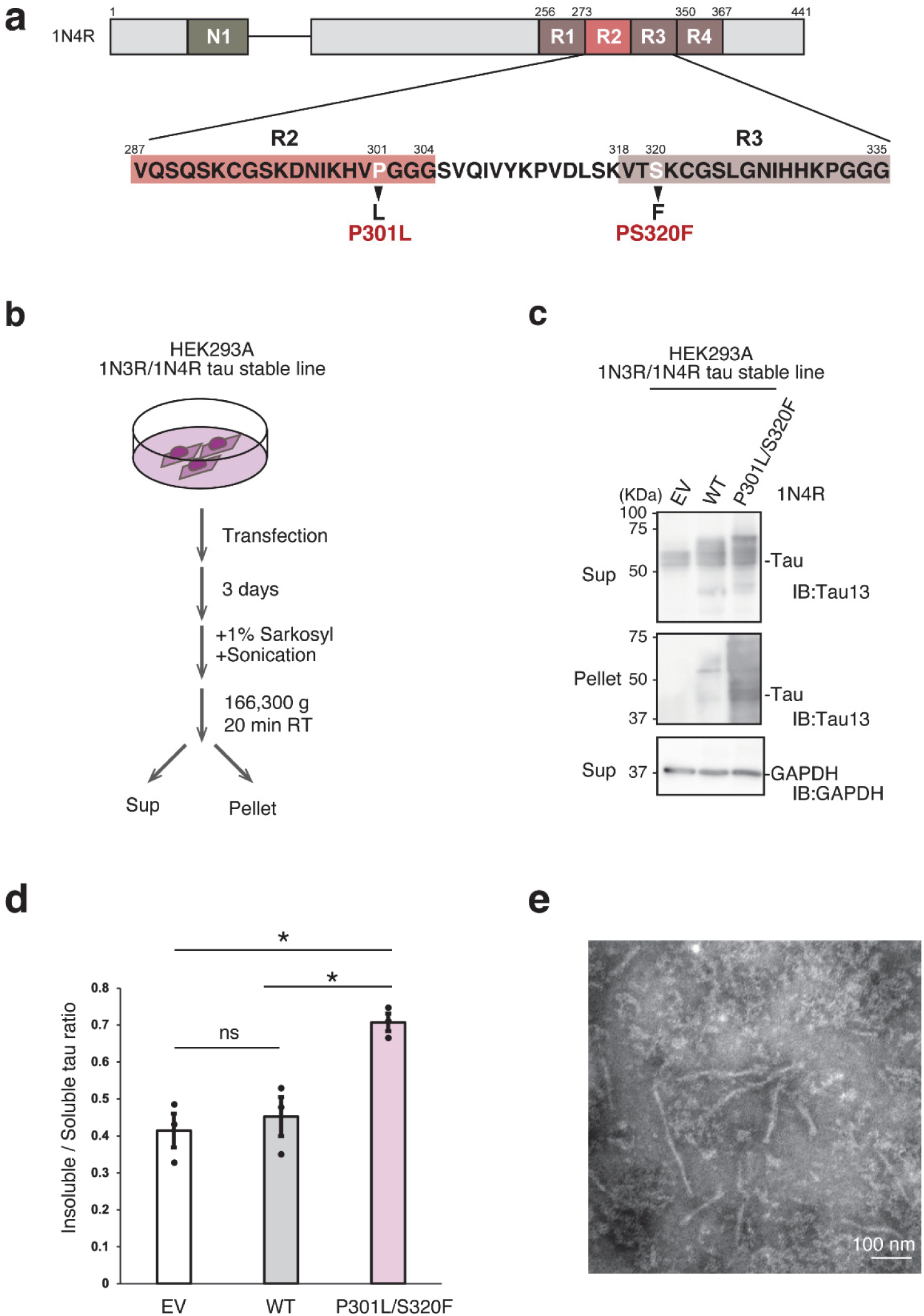
P301L/S320F double-mutant tau forms sarkosyl-insoluble tau filaments in cultured cells. **a.** Schematic representation of human 1N4R tau, showing the amino acid sequences of the microtubule-binding repeats (R1–R4), and the mutation sites P301L and S320F. **b.** Experimental design for assessing tau aggregation in HEK293A cells. Cells stably expressing 1N3R and 1N4R WT tau isoforms were transfected with plasmids encoding either 1N4R WT tau or 1N4R P301L/S320F double-mutant tau, followed by sarkosyl extraction to separate the soluble (Sup) and insoluble (Pellet) fractions. **c.** Western blot analysis of each fraction. Tau in the sarkosyl-insoluble fraction was prominently detected in the P301L/S320F group compared with the WT and empty-vector (EV) groups. GAPDH served as a loading control. **d.** Quantification of sarkosyl-insoluble tau normalized to soluble tau (mean ± SEM; n = 3 per group). Statistical significance was evaluated using a two-tailed Welch’s t-test. For all comparisons: p < 0.05 (*), ns = not significant. **e.** Resuspended sarkosyl-insoluble pellets obtained as in b were negatively stained with 2% phosphotungstic acid (pH 7.0) and observed under a transmission electron microscope. Scale bar is 100 nm.

### AAV-PHP.eB–mediated delivery of P301L/S320F double mutant tau induces intracellular tau aggregation in the mouse brain

In AD, tau aggregation is known to occur subsequent to the accumulation of Aβ. To model this coexisting Aβ and tau pathology *in vivo*, we employed *App*^NL-G-F^/*hMAPT* double knock-in mice [30,32]. This mouse line carries a knock-in human *APP* allele harboring the Swedish (KM670/671NL), Iberian (I716F), and Arctic (E693G) mutations (*App*^NL-G-F^), combined with a knock-in human *MAPT* (a.k.a., *hMAPT*) allele in which the endogenous mouse *Mapt* locus is replaced. In *App*^NL-G-F^ mice, age-dependent deposition of human Aβ begins around 2 months of age and reaches saturation by approximately 8 months. To induce tau aggregation under an Aβ-rich background, 9–11-month-old *App*^NL-G-F^/*hMAPT* mice were intravenously injected via the retro-orbital sinus with a recombinant AAV-PHP.eB expressing human P301L/S320F double-mutant tau under the human synapsin I promoter. Control groups included mice injected with WT human tau and non-injected mice (Fig. 2a). After two months, the brains were collected, hemisected, and used for immunohistochemical and biochemical analyses (Fig. 2a). AT8 immunoreactivity, a marker of pathological tau phosphorylation [60], revealed prominent AT8-positive signals in the CA2 region of the hippocampus and in layer II of the cerebral cortex in mice expressing P301L/S320F double-mutant tau (Fig. 2b). Quantification confirmed significant AT8 accumulation in the medial parietal association area (MPtA) (Fig. 2c, d) and hippocampal CA2 (Fig. 2e, f). Additionally, other antibodies detecting the phosphorylation-dependent tau pathology, such as AT180 [61], anti-pS396 [62], and PHF-1 [60], likewise detected exclusively in mice expressing P301L/S320F tau (Fig. S2– 4). Collectively, these findings indicate that P301L/S320F tau robustly induces the formation of phosphorylated tau aggregates *in vivo* without exogenous seeding.

**Figure 2.**
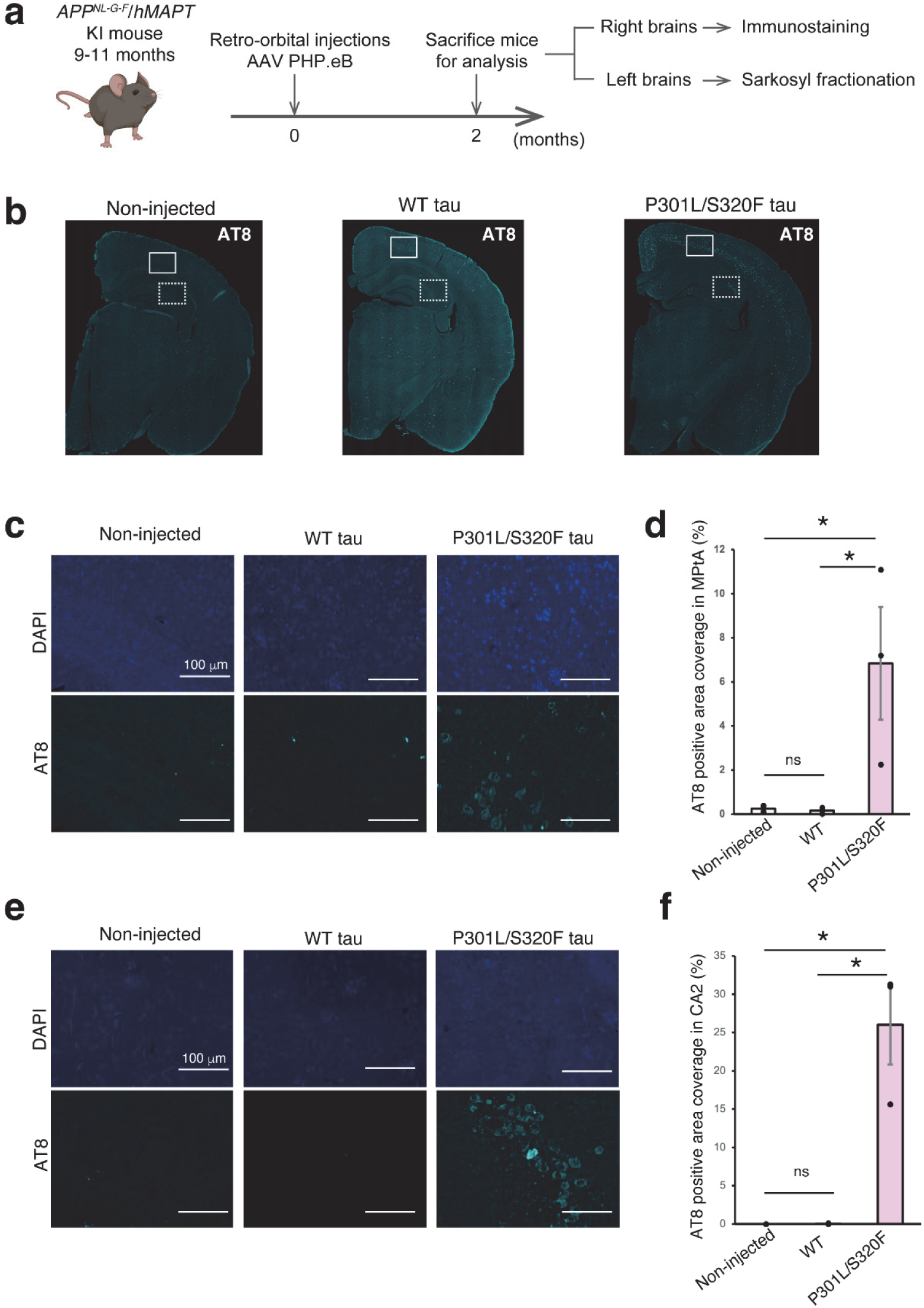
P301L/S320F tau induces phosphorylated tau aggregates in the brains of *App*^NL-G-F^/*hMAPT* mice. **a.** Experimental design. *App*^NL-G-F^/*hMAPT* double knock-in mice aged 9–11 months were intravenously injected via the retro-orbital sinus with the recombinant AAV-PHP.eB (2×10^11^vg/mouse). Brains were collected 2 months after injection for histological analysis. **b.** Immunostaining of brain hemispheres with AT8 antibody in mice expressing WT tau or P301L/S320F via AAV-PHP.eB, and from non-injected control mice. **c.** High-magnification images of the MPtA (boxed by a solid line) in b. Scale bars are 100 mm. **d.** Quantification of AT8-positive area in the MPtA. Statistical significance was determined by Welch’s two-tailed *t-*test. Mean ± SEM, n = 3. \**p* < 0.05, ns = not significant. **e.** High-magnification images of the hippocampal CA2 region in b. Scale bars are 100 mm. **f.** Quantification of AT8-positive area in the CA2 region. Statistical significance was determined by Welch’s two-tailed *t-*test. Mean ± SEM, n = 3. \**p* < 0.05, ns = not significant.

### Overexpressed P301L/S320F double-mutant tau in the mouse brain acts as a seed with strong aggregation propensity

We then biochemically analyzed the right hemisphere of the brains from *App*^NL-G-F^/*hMAPT* mice injected with AAV expressing P301L/S320F tau by sequential sarkosyl fractionation and western blotting (Fig. 3b). Quantification of the amount of the aggregated tau in the sarkosyl-insoluble fraction relative to the soluble fraction was markedly increased in mice expressing P301L/S320F double-mutant tau compared with those expressing WT tau (Fig. 3c). Notably, endogenous 3R as well as 4R tau were also detected within the insoluble fraction. Furthermore, murine endogenous tau was also detected in the sarkosyl-insoluble tau fraction in mice expressing P301L/S320F double-mutant tau (Fig. S5a.b). Pellets from the sarkosyl-insoluble fractions were analyzed by negative-stain electron microscopy using phosphotungstic acid and by immuno-electron microscopy (Fig. 3d). Only in the P301L/S320F group, fibrillar structures were observed that were immunoreactive for TauC4 [8], AT8, and anti-pS396 antibodies (Fig. 3e-g). Finally, we assessed the seeding activity of brain lysates using BS cells [40]. After transfection of the mouse brain homogenates, the proportion of FRET-positive cells was measured by flow cytometry only in the P301L/S320F group. In contrast, WT tau and uninjected controls showed no detectable FRET signal (Fig. S6). These data suggested that the double-mutant tau acted as a seed for endogenous tau protein regardless of repeat numbers or species.

**Figure 3.**
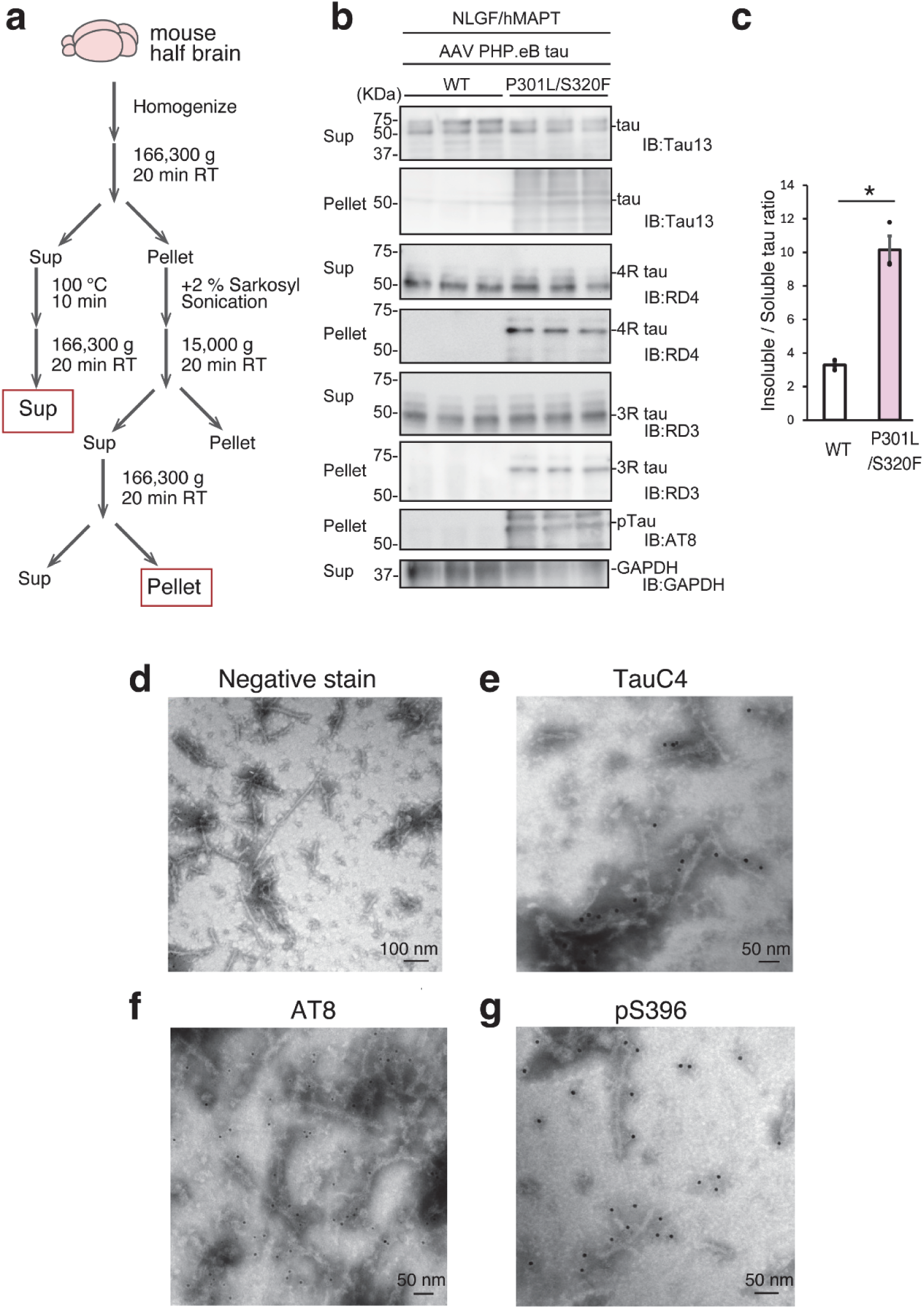
Evaluation of tau aggregation propensity and seeding activity in mouse brains expressing P301L/S320F tau. **a.** Experimental workflow. Following systemic administration of AAV-PHP.eB encoding either WT or P301L/S320F tau, mouse brain hemispheres were fractionated into soluble (Sup) and insoluble (Pellet) fractions by the sarkosyl extraction method. **b.** Western blot analysis of each fraction. **c.** Ratio of sarkosyl-insoluble to soluble tau (mean ± SEM; n = 3 per group), showing a significant increase in the P301L/S320F group (*p* < 0.05). Statistical significance was determined by Welch’s two-tailed *t-*test. **d.** Sarkosyl-insoluble pellets were negatively stained with 2% phosphotungstic acid (pH 7.0). **e**-**g**. Filaments positive for TauC4 (e), AT8 (f), and pS396 (g) antibodies were observed in the P301L/S320F group. Secondary antibodies used were 10 nm gold-conjugated anti-rabbit IgG for TauC4 and pS396 and 5 nm gold-conjugated anti-mouse IgG for AT8.

### Structural analysis of tau filaments derived from mice expressing P301L/S320F mutant tau by cryo-EM

To characterize the ultrastructure of tau assemblies formed *in vivo*, we analyzed the sarkosyl-insoluble tau filaments isolated from the mouse brain expressing the P301L/S320F double-mutant tau (PDB: 22JY) by cryo-EM (Fig. S7). The predominant filaments consisted of a single helical filament type (Fig. 4a, b), and the final reconstructed map reached a resolution of 2.24 Å. The filaments exhibited an approximate diameter of 7 nm and a helical pitch of ∼35 nm (Fig. 4a, b). The ordered core comprised residues 279–329 of tau, forming five β-strands connected by loops and turns (Fig. 4c, d). These β-strands corresponded to β1 (N279–S285), β2 (H299–L301), β3 (S305–Y310), β4 (V319–C322), and β5 (L325–H329). The five β-strands were stacked to form a layered β-sheet architecture that contributed to filament stabilization. The β-strands were linked by flexible loops, giving the overall structure a gently curved, folded conformation (Fig. 4c, d).

**Figure 4.**
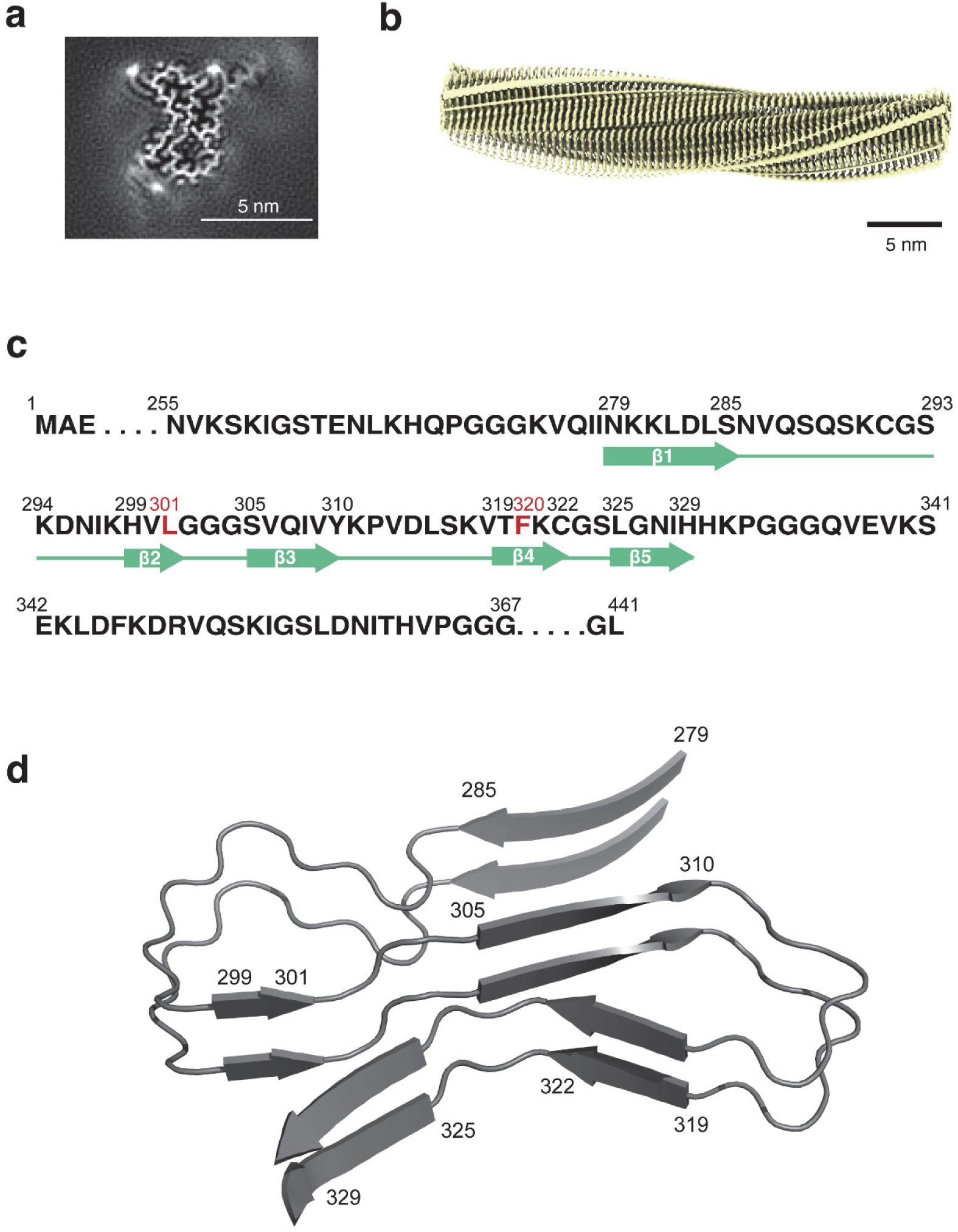
Cryo-EM structure of P301L/S320F tau filaments extracted from mouse brains. **a.** Representative cross-sections through the cryo-EM reconstruction of P301L/S320F tau filaments, shown perpendicular to the helical axis with a projected thickness of approximately one rung. The map resolution is 2.24 Å. Scale bar is 5 nm. **b.** Three-dimensional reconstruction of the P301L/S320F tau filaments obtained by helical reconstruction. Scale bar is 5 nm. **c.** Amino acid sequence of human 1N4R tau (residues 255–367) corresponding to the reconstructed region. The five β-strand regions (β1–β5) are indicated by green arrows. Mutation sites P301L and S320F are shown in red. **d.** Rendered view of the secondary structure elements in two successive rungs.

In addition, three prominent extra densities protruding from the filament surface were observed near lysine residues; although consistently positioned, their molecular identities remain unclear (Fig. 5a, b). The pathogenic substitutions, P301L and S320F, were each located within the second and fourth β-strands of the core, respectively. P301L formed hydrophobic interactions with I328 (Fig. 5c), whereas S320F was buried within a loop region, where its aromatic ring engaged in Z-axis π–π stacking that further stabilized the filament. The aromatic ring of S320F additionally established hydrophobic contacts with V309, P312, L315, and V318, likely increasing the stability of the entire loop structure. The Q307 side-chain amide groups (−CONH_2_) were positioned to form hydrogen bonds between adjacent layers aligned along the helical axis, further supporting inter-layer stabilization (Fig. 5d).

**Figure 5.**
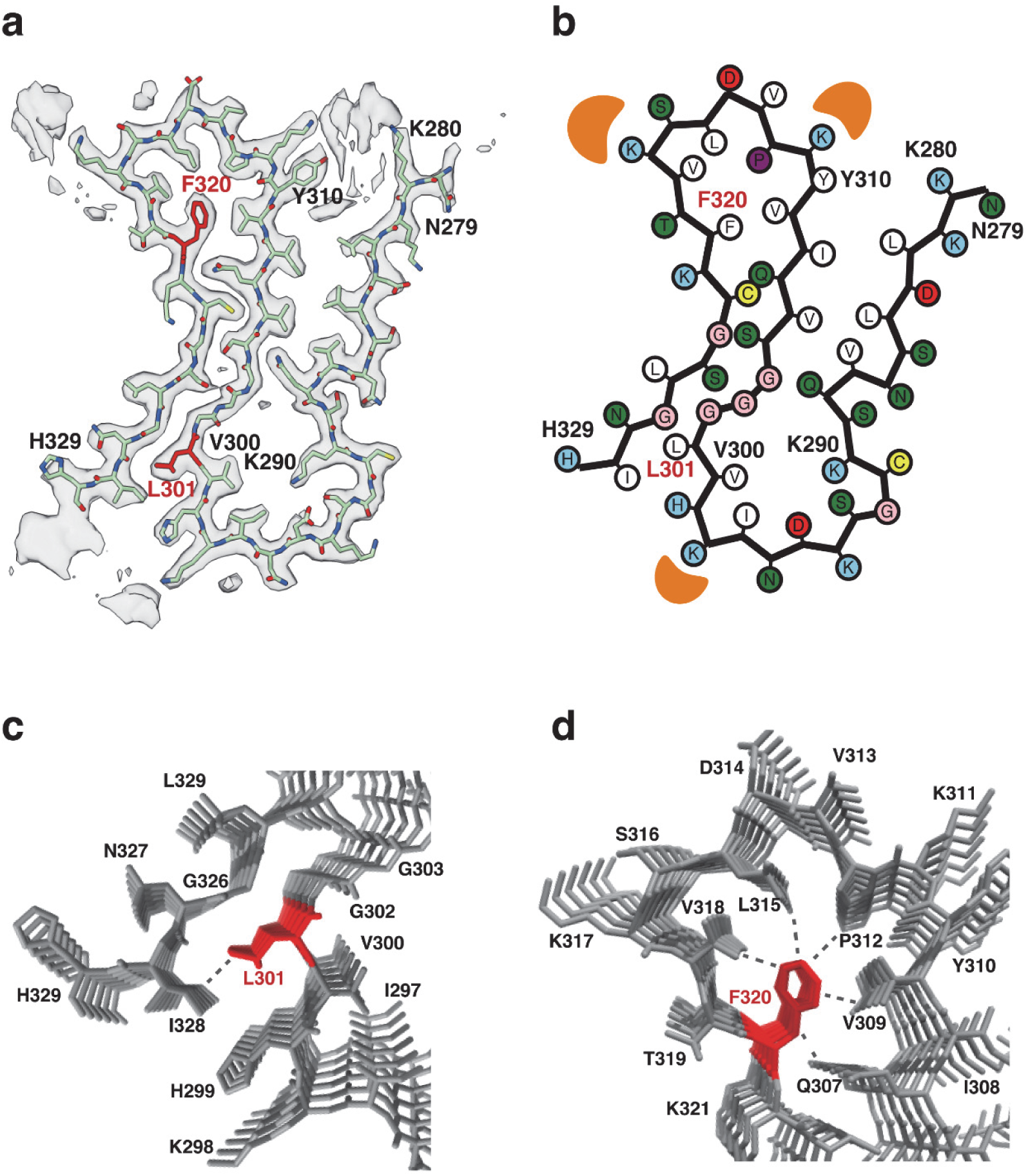
Local structural effects of P301L and S320F mutations on tau filaments. **a.** Sharpened cryo-EM density map of the P301L/S320F tau filament with the atomic model fitted. Major side chains are shown, and mutated residues L301 and F320 are highlighted in red. **b.** Schematic representation of the P301L/S320F tau filament fold. β-strands are indicated by thick lines with arrowheads. Orange semicircles indicate additional densities observed near the filament crossover region. **c.** Structure surrounding the P301L mutation site. The L301 residue (red) is positioned within the β-strand core and forms hydrophobic interactions with nearby residues, including V300 and I297. **d.** Structure surrounding the S320F mutation site. The F320 residue (red) is embedded within the loop region and forms hydrophobic interactions with V309, P312, L315, and V318, which likely contribute to the stabilization of the filament.

These hydrophobic interactions are not present in WT tau, where P301 and S320 are normally solvent-exposed. Therefore, the incorporation of these residues into the filament core represents a structural feature unique to the mutant filaments.

### The structure of P301L/S320F mutant tau filaments was distinct from previously reported patient-derived tau filament structures

To compare the structure of the P301L/S320F mutant tau filaments obtained in this study (magenta) with the structure of tau filaments derived from an FTDP-17 patient carrying the P301L mutation (PDB 9GG0; cyan) [63], we performed structural alignment. The results showed limited similarity, with an RMSD of 4.23 Å over 40 aligned residues (Fig. 6a). In the patient-derived fold, the filament bends at the P301L position, whereas in the P301L/S320F mutant, the filament bends at a different region. Moreover, in our structure, a hydrophobic interaction between P301L and I328—absent in the patient P301L fold—is observed. This is likely due to the S320F-induced folding of a unique loop through extensive hydrophobic interactions, bringing P301L into proximity with I328. The combined effects of P301L- and S320F-mediated interactions appear to rigidify the L301–H329 region, enabling the formation of a second compact loop in which V306 and I308 interlock with L282, L284, and V287 (Fig. 5a, b).

**Figure 6.**
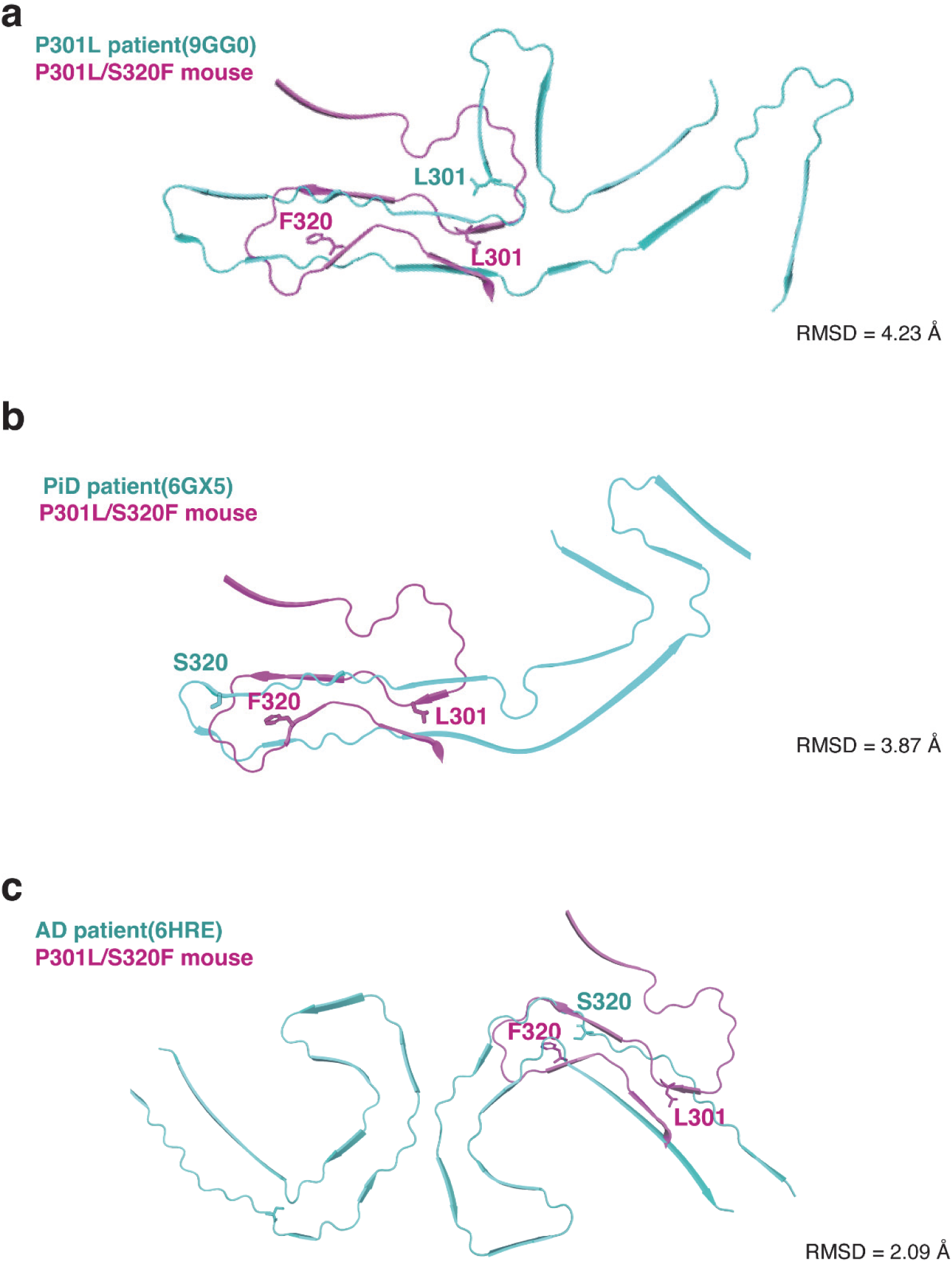
Comparison with previously reported patient-derived tau filament structures. **a.** Structural comparison between P301L/S320F mouse tau filaments (magenta) and P301L patient tau filaments (cyan, PDB: 9GG0), with an RMSD of 4.23 Å over 40 residues. **b.** Structural comparison between P301L/S320F mouse tau filaments (magenta) and PiD patient tau filaments (cyan, PDB: 6GX5), with an RMSD of 3.87 Å over 42 residues. **c.** Structural comparison between P301L/S320F mouse tau filaments (magenta) and AD patient tau filaments (cyan, PDB: 6HRE), with an RMSD of 2.09 Å over 16 residues.

In patients with FTDP-17 carrying the S320F mutation [25,26], Pick bodies composed of 3R tau are observed as a characteristic pathology [64]. We therefore compared our structure with the tau filament derived from PiD (PDB 6GX5; cyan)[65]. The residue numbering corresponds to the 441-aa human tau isoform, in which the R2 repeat (residues 275–305) is absent. The similarity was again low, with an RMSD of 3.87 Å over 42 residues (Fig. 6b).

Likewise, comparison with AD-derived PHFs (PDB 6HRE; cyan) [66] showed similarity only in a short loop near S320 (RMSD = 2.09 Å over 16 residues), whereas the overall fold was clearly distinct (Fig. 6c). Taken together, the fold obtained here is distinct from previously reported tau filament structures, including the Pick fold, the three-lobed fold, nor the Alzheimer PHF fold. Importantly, it also differs from the patient-derived P301L fold, suggesting that the combination of P301L and S320F mutations stabilizes a novel filament conformation not observed in any previously described human tauopathy.

## Discussion

P301L/S320F double-mutant has already been established as a rapidly developing tauopathy model characterized by aggregation of and accumulation of phosphorylated tau in neuronal cells, NFT pathology, cognitive impairment, synaptic protein abnormalities, and deficits in contextual memory [27]. In contrast, conventional tau transgenic mice typically require more than 6 months to develop detectable tau tangles. Therefore, AAV-mediated expression of P301L/S320F double-mutant tau can be regarded as a much more robust model of tau pathology. We further demonstrate that peripheral administration of the recombinant AAV-PHP.eB encoding P301L/S320F mutant tau rapidly induced the neuronal intracellular aggregation and filament formation in mouse brains without the injection of exogenous seeds to mouse brains. Owing to this combination of mutations, P301L/S320F double-mutant tau provides an excellent and not highly invasive tauopathy model in which tau pathology is markedly accelerated following AAV delivery.

Our cryo-EM analysis revealed that the extracted filaments consisted predominantly of single helical filaments, and the final reconstructed map reached a resolution of 2.24 Å. The ordered core was composed of tau residues 279–329, in which five β-sheets connected by loop and turn structures formed a compact three-folded architecture. This fold accounted for 55.9% of all tau filaments with clearly resolved β-sheets observed by cryo-EM. All filaments in which a defined helical pitch could be confidently determined exhibited the structure (Fig. 5). The remaining 44.1 % of filaments displayed a longer helical pitch or non-helical and disordered β-sheets, making structural determination difficult. The fold observed for P301L/S320F mutant tau in this study is distinct from the known tau folds reported in AD and PiD, nor from the structure derived from P301L patient brains (Fig. 6). This indicates that the combination of P301L and S320F generates a newly stabilized and unique three-dimensional conformation. Such structural insights provide important clues for understanding the diversity of tau filaments and the molecular basis of filament formation in tauopathies.

The S320F mutation is clinically associated with pathology resembling sporadic PiD, in which both straight and twisted tau filaments have been reported. However, whereas PiD inclusions are composed exclusively of 3R tau, the insoluble tau fraction from S320F carriers contains both 3R and 4R tau isoforms [25]. Although S320F carriers exhibit PiD-like straight filament formation and develop frontotemporal dementia, the coexistence of 3R and 4R tau suggests that the underlying filament architecture in these cases may differ from that of typical PiD, which is defined by the exclusive presence of 3R tau. Electron microscopy has further shown that the major tau species (∼80%) form straight filaments similar to those in AD [25,67]. *In vitro*, S320F tau has been shown to increase the rate of tau nucleation, shorten the lag phase of aggregation, and promote the formation of short protofibrils and oligomers [68,69], which may explain the ability of this mutant to aggregate in the absence of preformed seeds [70]. Taking these previous reports together with our structural data, S320F appears to form a stable loop through π– π stacking between F320 aromatic rings along the Z-axis and hydrophobic interactions with the surrounding residues V309, P312, L315, and V318. Because V309, P312, L315, and V318 are shared by both 3R and 4R tau, tau carrying the S320F mutation is likely to recruit both free 3R and 4R tau into the growing filaments. Thus, the loop structure centered on S320F may accelerate amyloid assembly and core formation, providing a mechanistic explanation for the presence of both 3R and 4R isoforms in the insoluble tau fractions from S320F carriers. Furthermore, comparison of the S320F-containing loop with the structure of tau filaments from PiD patients lacking this mutation reveals similarities in the length and packing of the major hairpin-like arm around residues 306–341 and in the packing of residues near position 320. Taken together, these observations suggest that the S320F mutation itself, rather than tau isoform composition alone, may contribute to the emergence of a PiD-like disease phenotype through its unique structural effects, even in filaments containing both 3R and 4R tau isoforms.

However, among filaments with a clearly defined pitch, we did not observe a robust fold that incorporated WT 3R or 4R tau. Such species likely belong to the population of filaments with excessively long pitch and partially ordered β-sheets in which structural determination was not feasible. This implies that, although S320F promotes filament formation through loop stabilization, it may not by itself be sufficient to generate highly ordered β-sheets within a short time frame in the model mouse. This interpretation is consistent with previous observations that AAV-mediated expression of S320F alone in neonatal ventricles does not produce substantial tau aggregation within three months [27].

In the present study, the combination of P301L and S320F mutations generated not only the S320F-dependent loop but also an additional hydrophobic interaction between L301 and I328. This hydrophobic contact appears to rigidify the base of the S320F-dependent loop, thereby defining a specific loop spanning 28 amino acids from L301 to I328. Formation of this L301–I328 loop exposes the hydrophobic residues V306 and I308 on the outer surface of the loop, allowing L282, L284, V287, and Q288 to pack against them and stabilize the three-folded architecture. Furthermore, interactions between N286 and S289 are likely to support and lock in this folding pattern. Collectively, these interactions contribute to the robust, compact three-folded structure observed for P301L/S320F double-mutant tau.

One of the key advantages of our model is the rapid induction of tau pathology. Whereas conventional models such as PS19 mice require more than 6 months for detectable tau aggregation, our system recapitulated tau accumulation and filament formation within 1–2 months. This should not be interpreted as a more faithful reproduction of human pathology, but rather as a powerful experimental system that enables rapid induction and analysis of tau aggregation. Because tau filament formation in our model was induced within at most 2 months in mouse brains, it is plausible that we captured a particularly stable and compact fold. WT 4R tau does not contain L301, and 3R tau lacks this region entirely; therefore, filaments that incorporate WT tau isoforms are expected to adopt structures distinct from the fold identified here, and their maturation may require longer periods. Long-term observation will thus be necessary to elucidate how WT tau is incorporated and how these filaments structurally evolve.

This study has several limitations: although peripheral administration of a recombinant AAV-PHP.eB achieved widespread gene delivery throughout the brain, transduction efficiency varied among regions, and the distribution of tau aggregates largely mirrored the pattern of AAV infection. In addition, because the number of transduced cells was limited, we did not observe large-scale neurodegeneration such as overt brain atrophy or extensive neuronal loss. Future studies combining region-specific intracranial injections with strategies to further enhance transduction efficiency may allow stronger pathology and more widespread neurodegeneration to be modeled.

Taken together, our findings demonstrate that P301L/S320F mutant tau promotes aggregation and filament formation in the absence of exogenous seeds, and that the resulting fold is clearly distinct from previously reported tau structures. These results deepen our mechanistic understanding of tau aggregation and provide a structural framework for exploring the basis of tauopathies and for identifying potential therapeutic targets.

## Conclusions

In this study, we established a non-invasive and rapid *in vivo* model of tau pathology by administering a recombinant AAV-PHP.eB from the periphery. Expression of the P301L/S320F double-mutant human tau in adult mouse brain induced robust tau aggregation without the need for exogenous seeds, accompanied by phosphorylation patterns commonly observed in AD, including AT8, AT180, pSer396, and PHF-1. The concomitant detection of early (AT8 and AT180) and late (pSer396 and PHF-1) phosphorylation epitopes indicates that tau phosphorylation progresses in a stepwise manner, leading to the accumulation of filamentous tau species that have reached an advanced pathological maturation stage. Our cryo-EM analysis of sarkosyl-insoluble tau filaments demonstrated that the combined P301L and S320F mutations cooperatively remodel the tau core to generate previously unreported filament fold distinct from known patient-derived structures. Together, these findings reveal the strong intrinsic aggregation propensity and unique fibrillization mechanism of P301L/S320F tau *in vivo*. The model presented here faithfully reproduces tau pathology on a background of humanized tau and Aβ deposition, providing a powerful platform for dissecting disease mechanisms and advancing therapeutic strategies for tauopathies.

## Abbreviations

AAV: adeno-associated virus
AD: Alzheimer disease
APP: amyloid precursor protein BS cells: biosensor cells
CA2: Cornu Ammonis 2
cryo-EM: cryo-electron microscopy
DAPI: 4’,6-diamidino-2-phenylindole
EM: electron microscopy
FRET: fluorescence resonance energy transfer
FTD: frontotemporal dementia
FTDP-17: frontotemporal dementia with Parkinsonism linked to chromosome 17
HEK293A: human embryonic kidney 293A
hMAPT: human microtubule-associated protein tau
MPtA: medial parietal association area
PBS: phosphate-buffered saline
RD: repeat domain
Sup: sarkosyl-soluble fraction Pellet: sarkosyl-insoluble fraction vg: viral genomes
WPRE: woodchuck hepatitis virus post-transcriptional regulatory element
WT: wild-type

## Declarations

### Ethics approval and consent to participate

All animal experiments were conducted in accordance with the guidelines of the Institutional Animal Care Committee of the Graduate School of Pharmaceutical Sciences at the University of Tokyo (protocol no. P3-20).

### Consent for publication

Not applicable.

### Availability of data and materials

Cryo-EM maps have been deposited in the Electron Microscopy Data Bank (EMDB) with the accession numbers EMD-68389 (P301L/S320F tau filament). Corresponding refined atomic models have been deposited in the Protein Data Bank (PDB) under accession numbers PDB: 22JY. The datasets used and analyzed during the current study are available from the corresponding author on reasonable request.

### Authors’ contributions

M.Ka performed cell and biochemical experiments, extracted filaments from mouse brains, analyzed the data, prepared figures, and wrote the original draft of the manuscript. T.K. supervised the biochemical experiments and provided technical guidance. H.Y. conducted cryo-EM imaging, performed structural reconstruction, and assisted in figure preparation. L.T. contributed to the optimization of experimental conditions for cell-based and mouse experiments. K.Y. performed immunohistochemical staining of mouse brain sections. A.S. carried out quantitative analyses of Western blot and immunohistochemical staining data. X.L. extracted tau filaments from cultured cells. S.T. constructed the P301L/S320F tau plasmid and established the AAV-PHP.eB systemic delivery protocol in mice. T.C.S. and T.S. established and provided *App^NL-G-F^/hMAPT* double knock-in mice. M.Ki supervised the structural analysis and provided critical input on the manuscript. T.T. conceived and supervised the overall project, secured funding, and served as the corresponding author.

## Acknowledgements

AAV production and purification were performed by the Department of Neurophysiology & Neural Repair, Gunma University Graduate School of Medicine, Gunma University (supported by JP21dm0207111). The authors are grateful to the Feinstein Institutes for Medical Research (Dr. Philippe Marambaud and Dr. Jeremy Koppel) for providing the PHF-1 antibody, originally developed by Dr. Peter Davies at Albert Einstein College of Medicine [71]. We acknowledge Viviana Gradinaru and Benjamin Deverman for their pioneering work in developing the AAV-PHP.eB capsid [42].

