## Supplementary information for "Cryo-EM structure of pro-aggregant P301L/S320F double-mutant tau filaments formed in mouse brains following peripheral AAV delivery"

### Author Information

[1]Laboratory of Neuropathology and Neuroscience, Graduate School of Pharmaceutical Sciences, The University of Tokyo, 7-3-1 Hongo, Bunkyo-ku, Tokyo 113-0033, Japan

[2]Department of Cell Biology and Anatomy, Graduate School of Medicine, The University of Tokyo, 7-3-1 Hongo, Bunkyo-ku, Tokyo 113-0033, Japan

[3]Laboratory for Proteolytic Neuroscience, RIKEN Center for Brain Science, 2-1 Hirosawa, Wako, Saitama 351-0198, Japan

[4]Department of Neurocognitive Science, Institute of Brain Science, Nagoya City University Graduate School of Medical Science, 1 Kawasumi, Mizuho-cho, Mizuho-ku, Nagoya, Aichi 467-8601, Japan

[#]These authors contributed equally to this work.

### Supplementary Figure Legends

**Figure S1. Structural features of the six human tau isoforms.**

This schematic representation illustrates the exon composition of the six tau isoforms (0N3R, 1N3R, 2N3R, 0N4R, 1N4R, 2N4R) generated from the *hMAPT* gene. The isoforms are defined by the presence or absence of the N-terminal insertions (N1, N2) and the microtubule-binding repeat domains (R1–R4).

**Figure S2. Peripheral administration of AAV-PHP.eB encoding P301L/S320F tau induces AT180-positive phosphorylated tau aggregates.**

**a**. Immunostaining of brain hemispheres with AT180 antibody in mice expressing WT tau or P301L/S320F via AAV-PHP.eB, and from non-injected control mice.

**b**. High-magnification images of the MPtA (boxed by solid line) in **a**. Scale bars are 100 µm.

**c**. Quantification of AT180 immunostaining in the MPtA. Statistical significance was determined by Welch's two-tailed *t*-test. Mean ± SEM, n = 3. $*p < 0.05$.

**d**. High-magnification images of the hippocampal CA2 region (boxed by dashed line) in **a**. Scale bars are 100 µm.

**e**. Quantification of AT180 immunoreactivity in the CA2 region. Statistical significance was determined by Welch's two-tailed *t*-test. Mean ± SEM, n = 3. $*p < 0.05$.

**Figure S3. Peripheral administration of AAV-PHP.eB encoding P301L/S320F tau induces pS396-positive phosphorylated tau aggregates.**

**a**. Immunostaining of brain hemispheres with pS396 antibody in mice expressing WT tau or P301L/S320F via AAV-PHP.eB and from non-injected control mice.

**b**. High-magnification images of the MPtA (boxed by solid line) in **a**. Scale bars are 100 μm.

**c**. High-magnification images of the hippocampal CA2 region (boxed by dashed line) in **a**. Scale bars are 100 μm.

**Figure S4. Peripheral administration of AAV-PHP.eB encoding P301L/S320F tau induces PHF-1-positive phosphorylated tau aggregates.**

**a**. Immunostaining of brain hemispheres with PHF-1 antibody in mice expressing WT tau or P301L/S320F via AAV-PHP.eB, and from non-injected control mice.

**b**. High-magnification images of the MPtA (boxed by solid line) in **a**. Scale bars are 100 μm.

**c**. High-magnification images of the hippocampal CA2 region (boxed by dashed line) in **a**. Scale bars are 100 μm.

**Figure S5. Peripheral administration of AAV-PHP.eB encoding 301L/S320F induces sarkosyl-insoluble tau in wild-type mice.**

**a**. Experimental timeline. Two-month-old wild-type C57BL/6J mice received an intravenous injection of a recombinant AAV-PHP.eB into the retro-orbital sinus. Brains were collected 2 months after injection for biochemical analysis.

**b**. AAV-PHP.eB–injected mice expressing either WT tau or P301L/S320F human tau, as well as non-injected control mice (non-injected), were processed using the sarkosyl extraction procedure described in **Fig. 3a**. These fractions were analyzed by western blotting. Total tau was detected using the anti-tau antibody Tau13, and GAPDH was used as a loading control.

**Figure S6. FRET biosensor assay of P301L/S320F tau induced to mouse brains.**

**a**. Representative FACS histograms of BS cells incubated with mouse brain homogenates. FRET-positive signals were displayed as yellow events within the blue triangular gating region. FRET-positive cells were detected using flow cytometry with Brilliant Violet 510 / Brilliant Violet 421.

**b-d**, Representative flow cytometry plots of BS cells without homogenate (**b**), BS cells incubated with brain homogenate from *App*$^{NL\text{-}G\text{-}F}$/*hMAPT* mice expressing WT tau (**c**), and BS cells incubated with brain homogenate from *App*$^{NL\text{-}G\text{-}F}$/*hMAPT* mice expressing P301L/S320F human tau (**d**).

**e**, Quantification of the proportion of FRET-positive cells (mean ± SEM; n = 3 per group). Statistical significance was evaluated using a two-tailed Welch's *t*-test. For all comparisons: $p < 0.05$ (*), $p < 0.01$ (**), *ns* = not significant.

**Figure S7. Summary of the image processing of P301L/S320F human tau filaments**

Top: Cryo-EM pipeline for P301L/S320F human tau filaments.

Bottom left: Local resolution map, colored from blue (high resolution) to red (lower resolution).

Bottom right: FSC curves, demonstrating a resolution of 2.24 Å. FSC curves reflect unmasked (green), masked (blue), and corrected (black) estimates of map resolution.

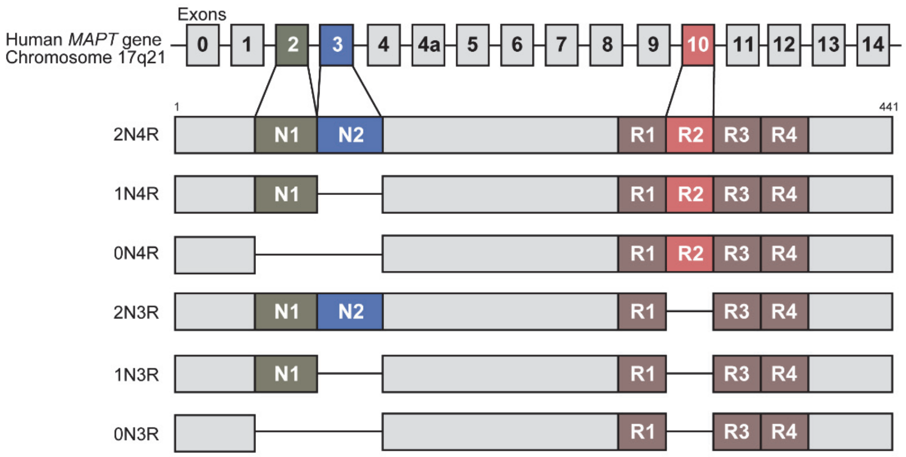
Exons
Human *MAPT* gene
Chromosome 17q21
0
1
2
3
4
4a
1
441
2N4R
1N4R
0N4R
2N3R
1N3R
0N3R
N1
N2
R1
R2
R3
R4

**Fig, S1**

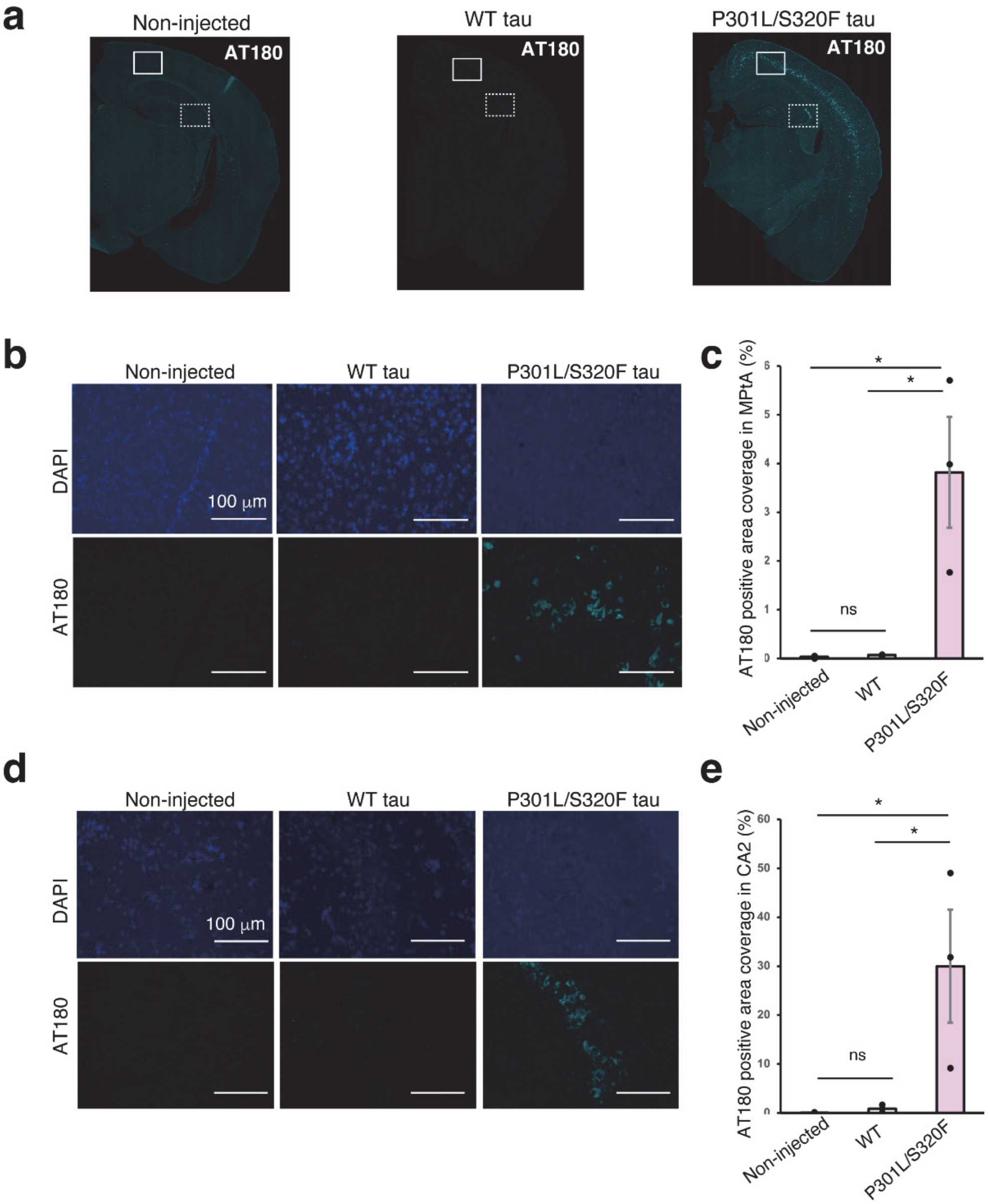
a
Non-injected
WT tau
P301L/S320F tau
AT180
b
Non-injected
WT tau
P301L/S320F tau
DAPI
AT180
100 μm
c
AT180 positive area coverage in MPtA (%)
ns
Non-injected
WT
P301L/S320F
d
Non-injected
WT tau
P301L/S320F tau
DAPI
AT180
100 μm
e
AT180 positive area coverage in CA2 (%)
ns
Non-injected
WT
P301L/S320F

**Fig, S2**

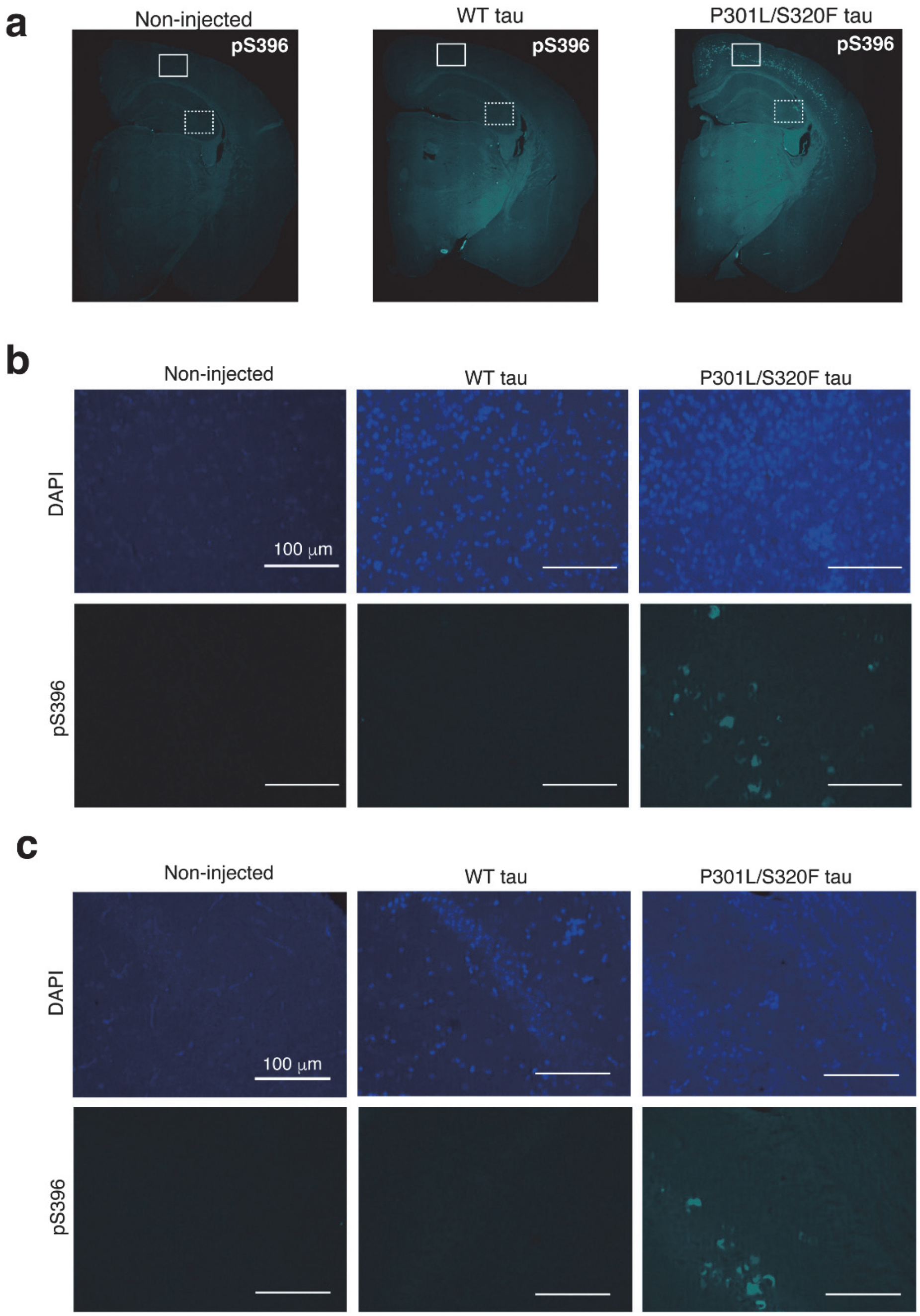
a
Non-injected
WT tau
P301L/S320F tau
pS396
pS396
pS396
b
Non-injected
WT tau
P301L/S320F tau
DAPI
100 µm
pS396
c
Non-injected
WT tau
P301L/S320F tau
DAPI
100 µm
pS396

**Fig, S3**

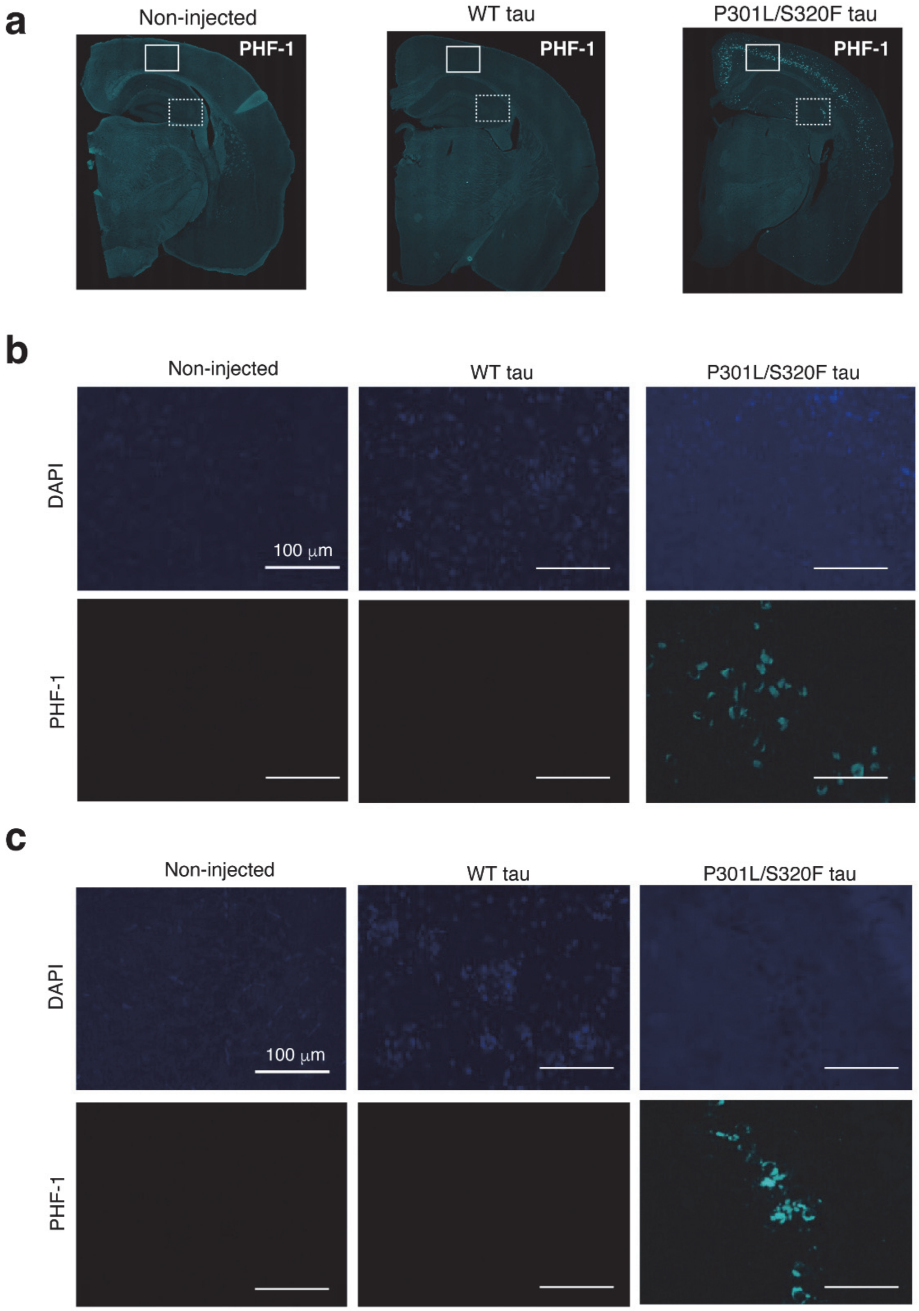
a
Non-injected
WT tau
P301L/S320F tau
PHF-1
PHF-1
PHF-1
b
Non-injected
WT tau
P301L/S320F tau
DAPI
PHF-1
100 μm
c
Non-injected
WT tau
P301L/S320F tau
DAPI
PHF-1
100 μm

**Fig, S4**

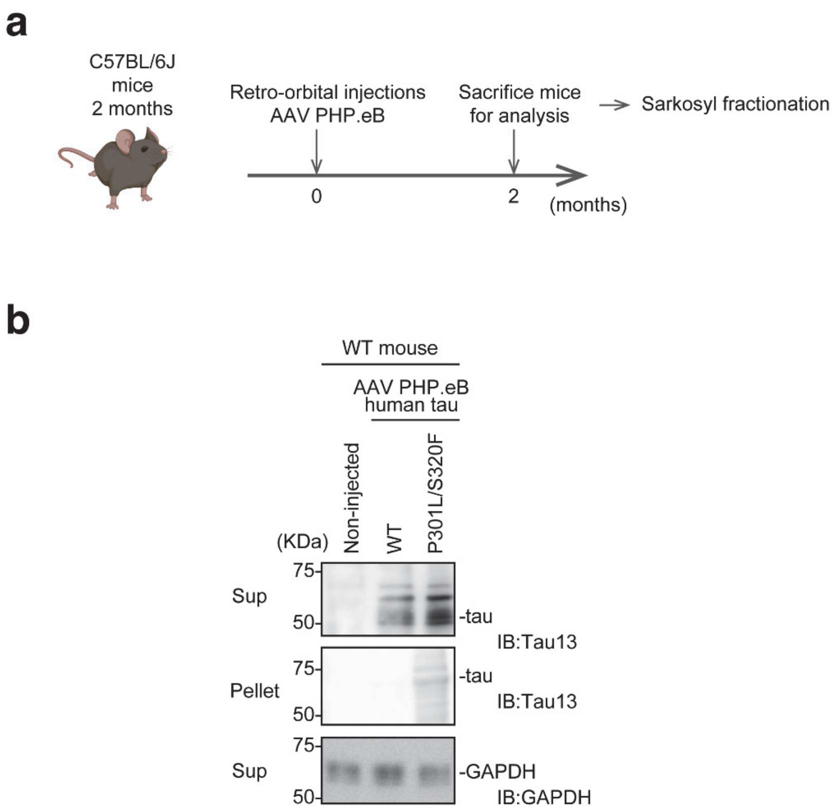
a
C57BL/6J
mice
2 months
Retro-orbital injections
AAV PHP.eB
Sacrifice mice
for analysis
Sarkosyl fractionation
0
2
(months)
b
WT mouse
AAV PHP.eB
human tau
Non-injected
WT
P301L/S320F
(KDa)
75
50
Sup
-tau
IB:Tau13
Pellet
-tau
IB:Tau13
Sup
-GAPDH
IB:GAPDH

**Fig, S5**

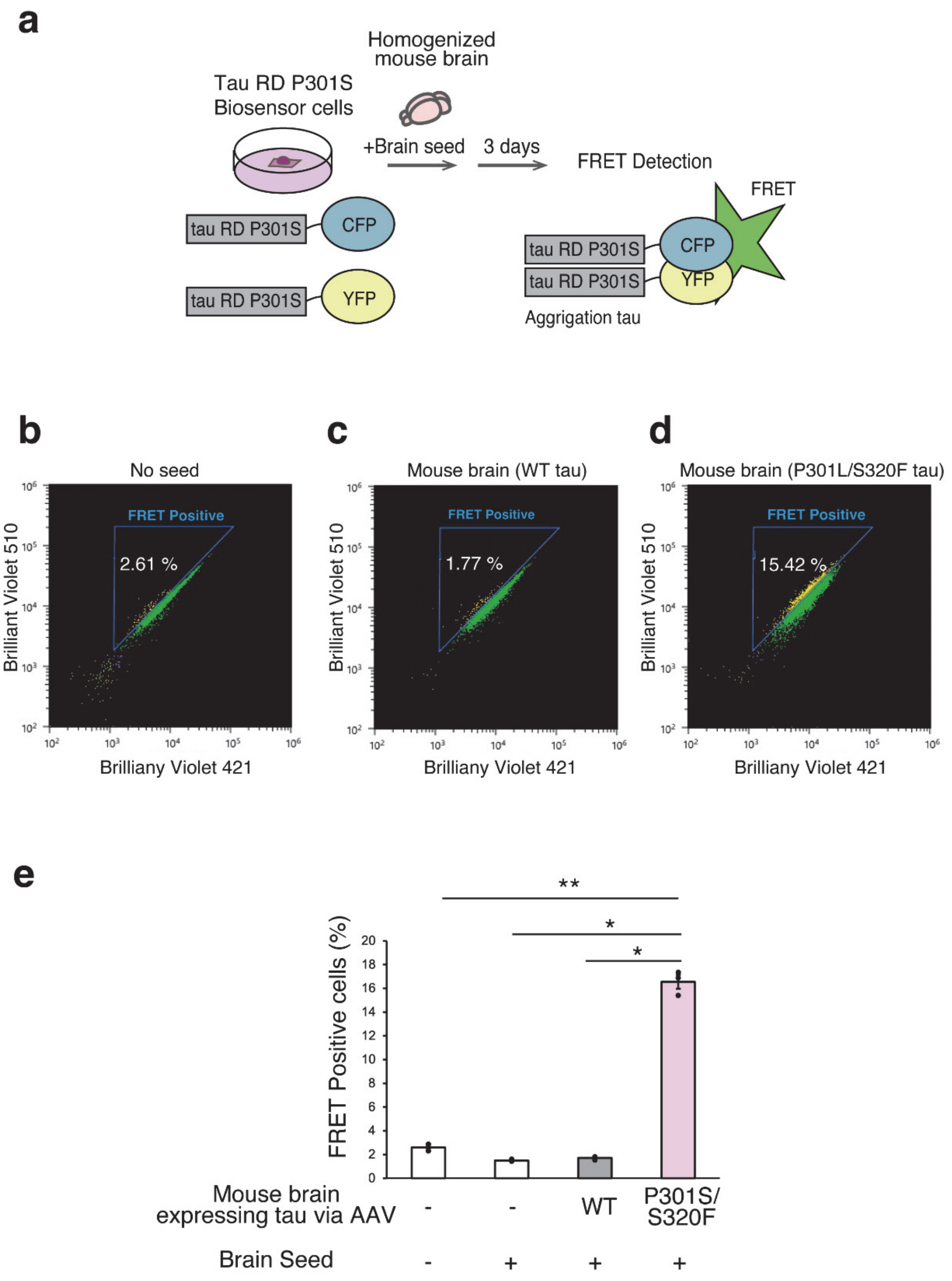

109

110 **Fig, S6**

111

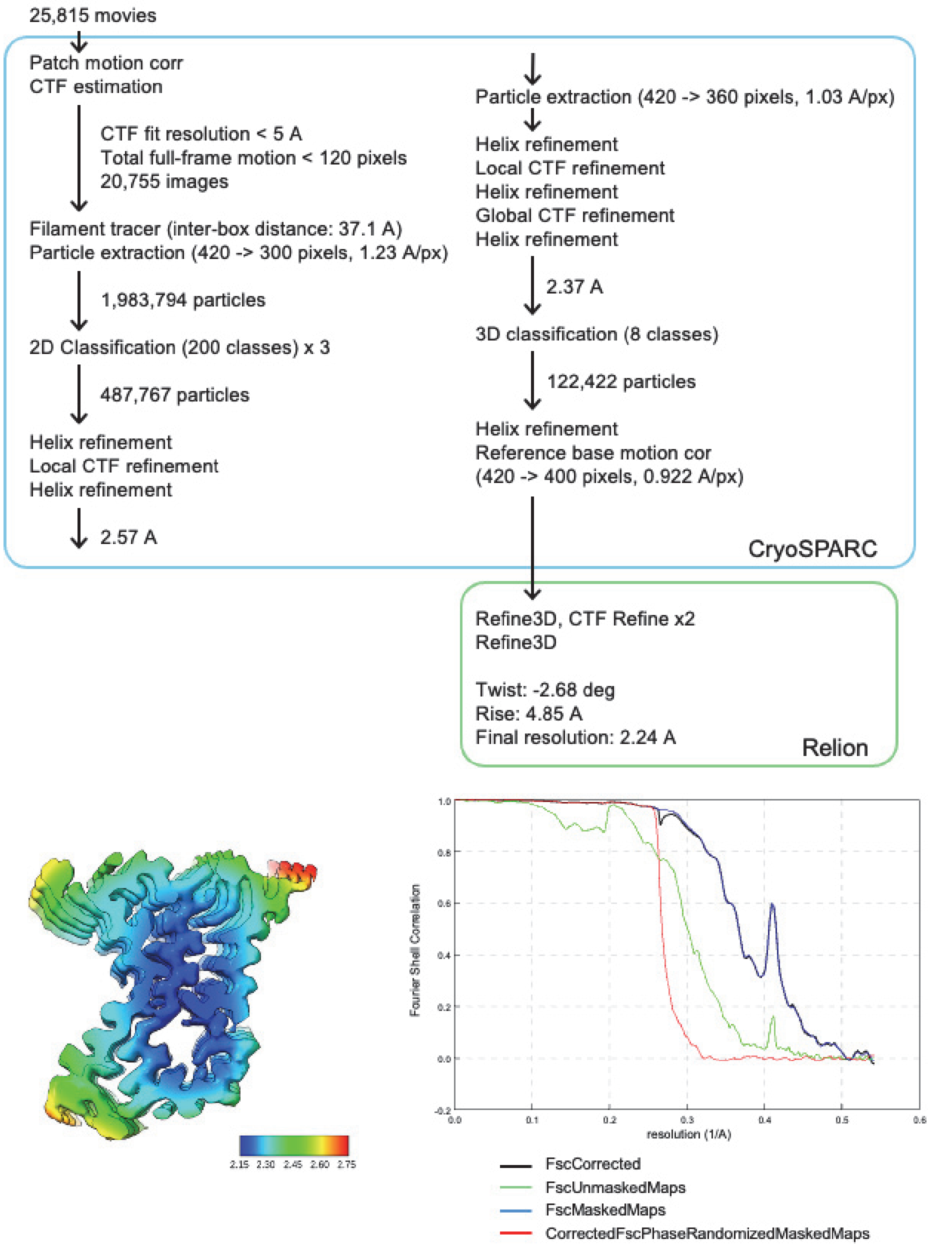
25,815 movies
Patch motion corr
CTF estimation
CTF fit resolution < 5 A
Total full-frame motion < 120 pixels
20,755 images
Filament tracer (inter-box distance: 37.1 A)
Particle extraction (420 -> 300 pixels, 1.23 A/px)
1,983,794 particles
2D Classification (200 classes) x 3
487,767 particles
Helix refinement
Local CTF refinement
Helix refinement
2.57 A
Particle extraction (420 -> 360 pixels, 1.03 A/px)
Helix refinement
Local CTF refinement
Helix refinement
Global CTF refinement
Helix refinement
2.37 A
3D classification (8 classes)
122,422 particles
Helix refinement
Reference base motion cor
(420 -> 400 pixels, 0.922 A/px)
CryoSPARC
Refine3D, CTF Refine x2
Refine3D
Twist: -2.68 deg
Rise: 4.85 A
Final resolution: 2.24 A
Relion
2.15 2.30 2.45 2.60 2.75
Fourier Shell Correlation
resolution (1/A)
FscCorrected
FscUnmaskedMaps
FscMaskedMaps
CorrectedFscPhaseRandomizedMaskedMaps

**Fig, S7**
